# Aldehyde inactivation of the RicR regulon sensitizes *Mycobacterium tuberculosis* to copper

**DOI:** 10.1101/2022.09.30.510424

**Authors:** Gina Limón, Nora M. Samhadaneh, Alejandro Pironti, K. Heran Darwin

**Affiliations:** Department of Microbiology, New York University Grossman School of Medicine, 430 E. 29^th^ Street, New York, NY, 10016, USA; Antimicrobial-Resistant Pathogens Program, New York University Grossman School of Medicine, 430 E. 29^th^ Street, New York, NY, 10016, USA; Microbial Computational Genomic Core Lab, New York University Grossman School of Medicine, 430 E. 29^th^ Street, New York, NY, 10016, USA

## Abstract

*Mycobacterium tuberculosis* is a major human pathogen and the causative agent of tuberculosis disease. While *M. tuberculosis* can persist in the presence of host-derived antimicrobials like nitric oxide and copper, bacteria defective for proteasome activity are highly sensitive to these molecules, making the proteasome an attractive target for drug development. Previous work linked nitric oxide susceptibility with the accumulation of at least one aldehyde in an *M. tuberculosis* mutant lacking proteasomal degradation. In this study, we show that this aldehyde accumulation is also responsible for copper sensitivity in this strain. Furthermore, we show the exogenous addition of aldehydes to wild-type *M. tuberculosis* cultures sensitizes bacteria to copper. We determined that aldehydes directly affect the activity of two members of the RicR (<u>r</u>egulated <u>i</u>n <u>c</u>opper <u>r</u>epressor) regulon, resulting in the reduced production and function of critical copper-responsive proteins. This study is the first to mechanistically describe how aldehydes can render *M. tuberculosis* susceptible to an existing host defense, and could support a broader role for aldehydes in controlling *M. tuberculosis* infections.

**IMPORTANCE:** *M. tuberculosis* is a leading cause of death by a single infectious agent, causing 1.5 million deaths annually. An effective vaccine for *M. tuberculosis* infections is currently lacking, and prior infection does not typically provide robust immunity to subsequent infections. Nonetheless, immunocompetent humans can control *M. tuberculosis* infections for decades. For these reasons, a clear understanding of how mammalian immunity inhibits mycobacterial growth is warranted. In this study, we show aldehydes can increase *M. tuberculosis* susceptibility to copper. Given that activated macrophages produce increased amounts of aldehydes during infection, we propose host-derived aldehydes target critical bacterial survival pathways, making aldehydes a previously unappreciated antimicrobial defense.

## INTRODUCTION

*Mycobacterium tuberculosis* is a major human pathogen and the causative agent of tuberculosis disease (TB). TB is responsible for more than 1 million deaths annually and was the deadliest infectious disease worldwide prior to the SARS-CoV-2 pandemic (https://www.who.int/news-room/fact-sheets/detail/tuberculosis). TB can be cured by lengthy treatments with multiple antibiotics; however, antibiotic-resistant strains of *M. tuberculosis* are increasingly prevalent. Thus, an improved understanding of *M. tuberculosis* pathogenesis and existing host responses to the bacteria are required to aid in the development of new treatment strategies.

Macrophages, a primary niche of *M. tuberculosis*, require the production of nitric oxide (NO) for robust resistance to various infections, in particular *M. tuberculosis* (1, 2). NO is a free radical that can form toxic reactive intermediates that damage molecules including nucleic acids, proteins, and lipids [reviewed in (1)]. While the precise mechanism of NO-mediated toxicity to *M. tuberculosis* is unknown, *M. tuberculosis* requires a Pup-proteasome system (PPS) to resist NO (3). In the PPS, numerous proteins that are destined for degradation are post-translationally modified with prokaryotic ubiquitin-like protein (Pup) by the ligase PafA (proteasome accessory factor A) [reviewed in (4)]. Pup acts as a signal that directly binds to a hexameric ATPase, Mpa (also known as ARC in non-mycobacteria), that unfolds pupylated proteins and delivers them into a proteasome core protease. In the absence of either Mpa or PafA, numerous proteins fail to be degraded. When the PPS is nonfunctional in *M. tuberculosis*, the enzyme lonely guy (Log) accumulates (5). Log plays a role in the production of adenine-based hormones called cytokinins, which in plants are required for normal growth and development (6). *M. tuberculosis* is not found in plants and instead uses cytokinins to induce gene transcription to alter the mycobacterial cell envelope (5, 7). In plants, cytokinins are enzymatically broken down into adenine and various aldehydes (8). While it is unknown how cytokinins are metabolized in *M. tuberculosis*, at least one cytokinin-associated aldehyde, *para*-hydroxybenzaldehyde (*p*HBA), measurably accumulates in a PPS-defective strain of *M. tuberculosis* and contributes to NO sensitivity (5).

In addition to NO, there is evidence that macrophages use copper (Cu) to defend against *M. tuberculosis* and other bacterial infections (9-13). *M. tuberculosis* possesses two defined Cu-responsive systems to mitigate Cu toxicity: the CsoR (<u>c</u>opper-<u>s</u>ensitive <u>o</u>peron <u>r</u>epressor) operon (14) and the RicR (<u>r</u>egulated in <u>c</u>opper <u>r</u>epressor) regulon (15). CsoR and RicR are paralogues that contribute to Cu resistance (12, 13, 15-17), although the RicR regulon plays a more substantial role in Cu resistance *in vitro* and in virulence in mice (12, 17). When Cu concentrations are low, RicR binds to the promoters of five loci, including the *ricR* promoter, preventing their expression. When Cu levels increase, Cu binds to RicR, preventing it from associating with DNA and allowing gene expression. Two RicR regulon gene products, MmcO, a multicopper oxidase, and MymT, a Cu-metallothionein, confer *M. tuberculosis* resistance to Cu; deletion of the genes encoding these proteins results in hyper-susceptibility to Cu *in vitro* (12, 16, 18). While deletion of any single RicR-regulated gene does not attenuate *M. tuberculosis* growth in C57BL6/J mice, the production of a “Cu-blind” RicR protein that constitutively represses expression of the entire regulon results in highly Cu-sensitive bacteria that are attenuated for growth in mice (12).

Our lab reported that *M. tuberculosis* PPS mutants are hypersensitive to Cu (12), which we reason is due to the repression of the RicR regulon in these strains (15). Given that aldehyde accumulation in PPS mutants sensitizes *M. tuberculosis* to NO (5), we hypothesized that aldehydes also cause Cu sensitivity. In this work, we determined that in a PPS-deficient *M. tuberculosis* strain, a mutation that prevents the accumulation of *p*HBA suppresses its Cu-sensitive phenotype. Additionally, we found *p*HBA can disrupt the production and activities of two Cu-binding proteins. Finally, we showed that an aldehyde that accumulates in macrophages during *M. tuberculosis* infections, methylglyoxal (MG), can increase *M. tuberculosis* susceptibility to Cu, suggesting host-derived aldehydes have the potential to inactivate Cu resistance *in vivo*. Collectively, our studies suggest that aldehydes control *M. tuberculosis* infections, partly by rendering bacteria hypersensitive to Cu.

## RESULTS

### Cu sensitivity of a proteasomal degradation-deficient *M. tuberculosis* strain is suppressed by disruption of *log*

*M. tuberculosis* strains with disruptions in *mpa* or *pafA* are hypersensitive to Cu (12). We previously showed that an *mpa* mutant is hypersensitive to NO due to the accumulation of the PPS substrate Log; a transposon insertion in *log* in an *mpa* strain suppresses its NO-sensitive phenotype (5). We tested if this same mutation could suppress the Cu-sensitive phenotype of a PPS mutant. We confirmed a Δ*mpa::hyg* (“*mpa*”) mutant is highly sensitive to Cu. We also found that transposon disruptions in *log* (*log*::MycoMarT7, “*log*”) in the *mpa* strain restored WT Cu resistance (**Fig. 1A**). This result suggested one or more accumulated aldehydes in a PPS mutant contributes its Cu-sensitive phenotype.

**Fig. 1.**
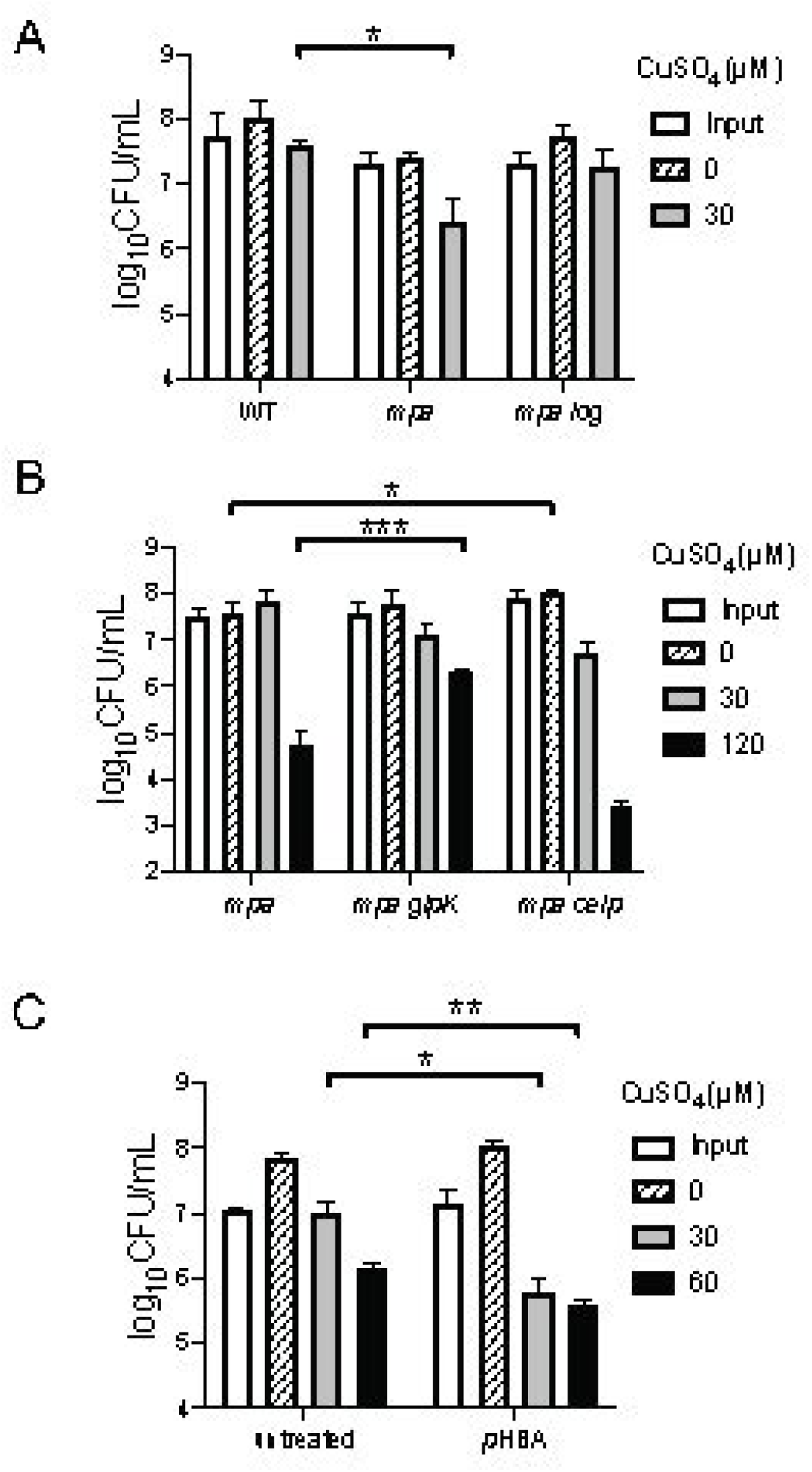
Aldehydes sensitize *M. tuberculosis* to copper. **(A)** Survival of *M. tuberculosis* strains (x-axis) in various concentrations of CuSO_4_ for 10 days. CFU were enumerated after 14 to 21 days of incubation on solid 7H11c medium. **(B)** Cu sensitivity assay performed as in (A) with *mpa* NO-sensitivity suppressor strains. **(C)** Cu sensitivity assays performed with WT *M. tuberculosis* incubated with or without 1.2 mM *p*HBA for 24 hours in Sauton media prior to treatment with CuSO_4_, as in (A). Data are representative of two independent experiments, each done in technical triplicate. All quantifications were analyzed for significance using an unpaired t-test with *P* < 0.05.

Previous work from our lab found two additional mutations that suppressed the NO sensitivity of an *mpa* mutant (5). These suppressor mutations are in *glpK*, which encodes glycerol kinase, and in the promoter for *cei* (Rv2700), which encodes a possible secreted alanine rich protein involved in cell membrane integrity (19). We tested Cu sensitivity of these strains and found that the *glpK*::MycoMarT7 mutation increased Cu resistance relative to the parental *mpa* strain (**Fig. 1B**, *mpa glpK*). In contrast, the transposon insertion in the *cei* promoter (*mpa ceip*) did not restore Cu resistance to parental levels (**Fig. 1B**, *mpa ceip*).

We previously showed that the addition of *p*HBA, an aldehyde that accumulates in an *mpa* mutant, sensitizes WT *M. tuberculosis* to NO (5), thus, we tested if exogenously added *p*HBA could sensitize *M. tuberculosis* to Cu. We used 1.2 mM *p*HBA for all experiments in this study given that this concentration is non-toxic on its own but can robustly synergize with NO to sterilize *M. tuberculosis* cultures (5). We attempted to add *p*HBA and Cu simultaneously to *M. tuberculosis* cultures; however, the presence of exogenous *p*HBA appeared to inhibit Cu-dependent killing (data not shown). To circumvent this problem, we reasoned that preincubation of bacteria with *p*HBA prior to Cu exposure would allow for changes that affect Cu resistance. We therefore preincubated WT *M. tuberculosis* with *p*HBA for 24 hours before exposing the bacteria to Cu. As previously reported, *p*HBA alone had no effect on CFUs recovered (**Fig. 1C**) (11) but we found incubating bacteria with *p*HBA before Cu exposure significantly reduced the number of colony-forming units (CFUs) recovered compared to Cu treatment alone (**Fig. 1C**). This result supported our hypothesis that the presence of *p*HBA sensitizes *M. tuberculosis* to Cu.

### Expression of the RicR regulon is reduced in *p*HBA-treated *M. tuberculosis*

To investigate the basis of Cu sensitivity in *p*HBA-treated *M. tuberculosis*, we performed RNA-Seq on WT *M. tuberculosis* treated with *p*HBA (**Table S1**, see **Table 1** for strains). We observed a majority of genes that were differentially regulated in *p*HBA-treated WT bacteria showed similar patterns of gene expression previously observed in PPS *M. tuberculosis* mutants (15) (**Table 2** and **Table 3**, gray rows). The substantial overlap between the *p*HBA-treated and PPS-mutant transcriptomes suggested that one or more aldehydes were responsible for the transcriptional phenotypes observed in PPS mutants. The shared differentially regulated genes included all eight members of the Cu-sensing RicR regulon, which were down-regulated (**Table 2**). As described previously, RicR is a repressor that dissociates from DNA in the presence of Cu, leading to the expression of genes from five different promoters (15). The CsoR operon also showed this pattern of repression in PPS mutant strains and *p*HBA-treated *M. tuberculosis* (**Table 2**). Like RicR, CsoR releases repression of its operon after binding Cu to help *M. tuberculosis* maintain Cu homeostasis (14, 20). For example, the CsoR-regulated gene *ctpV* encodes a cation transporter implicated in Cu resistance and virulence (17).

**Table 1.**
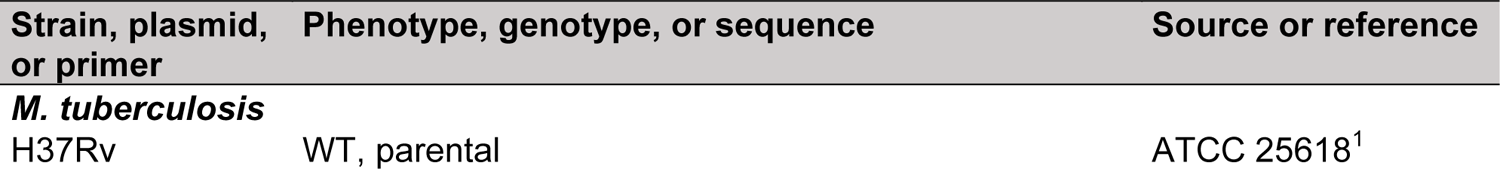
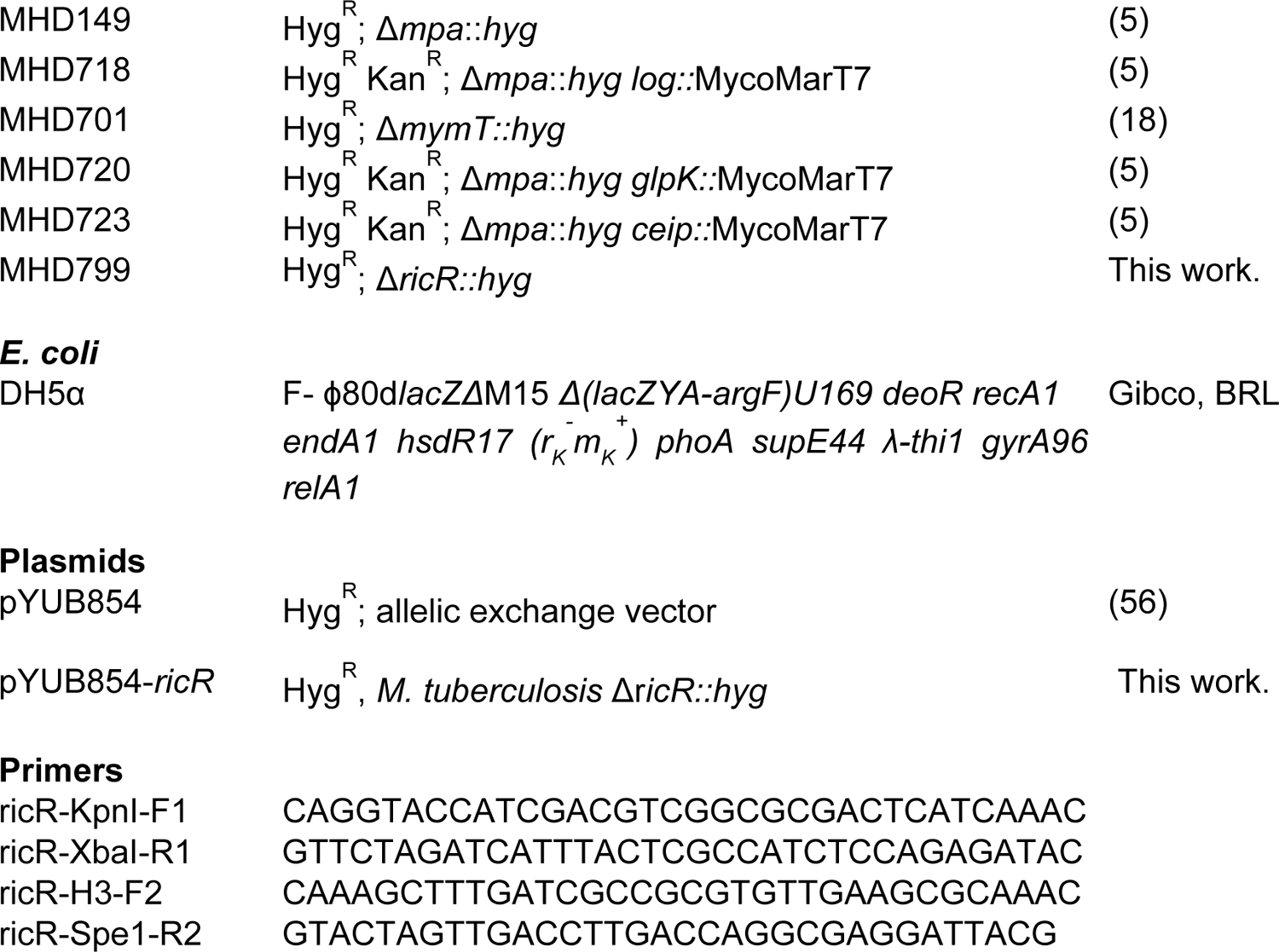
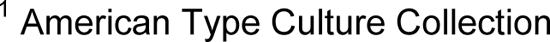
Strains, plasmids, and primers used in this study.

**Table 2.**
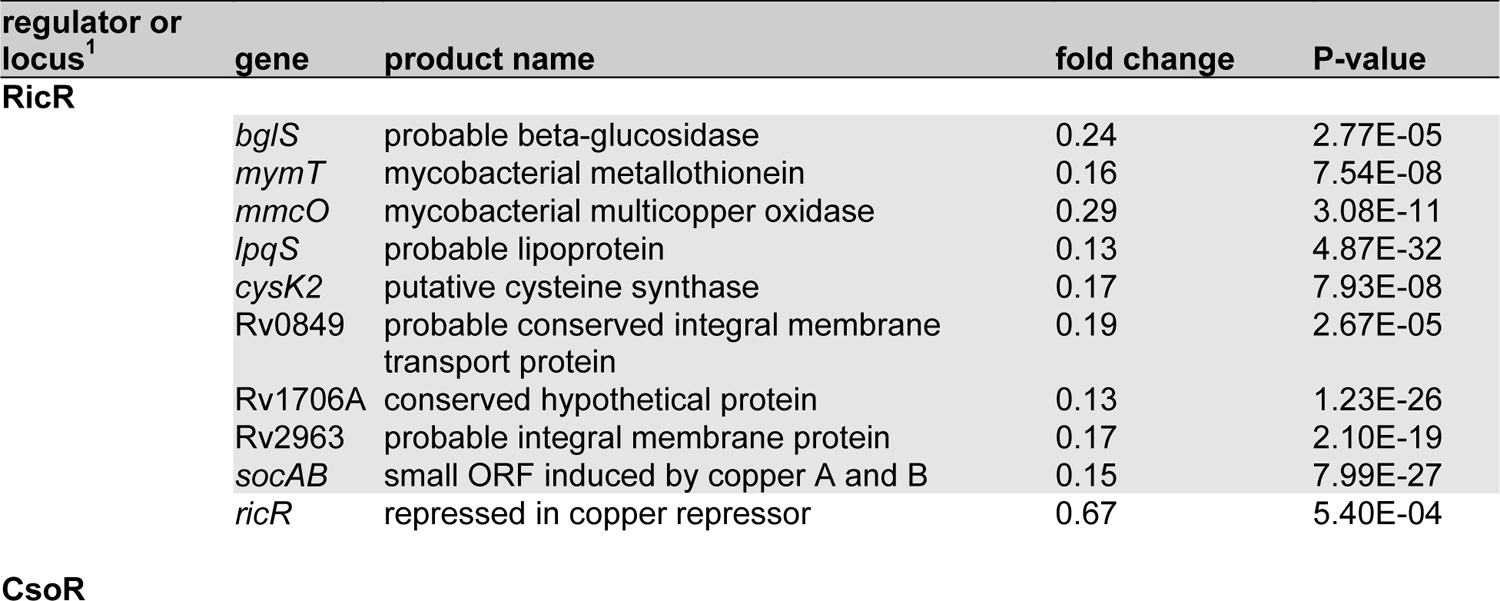
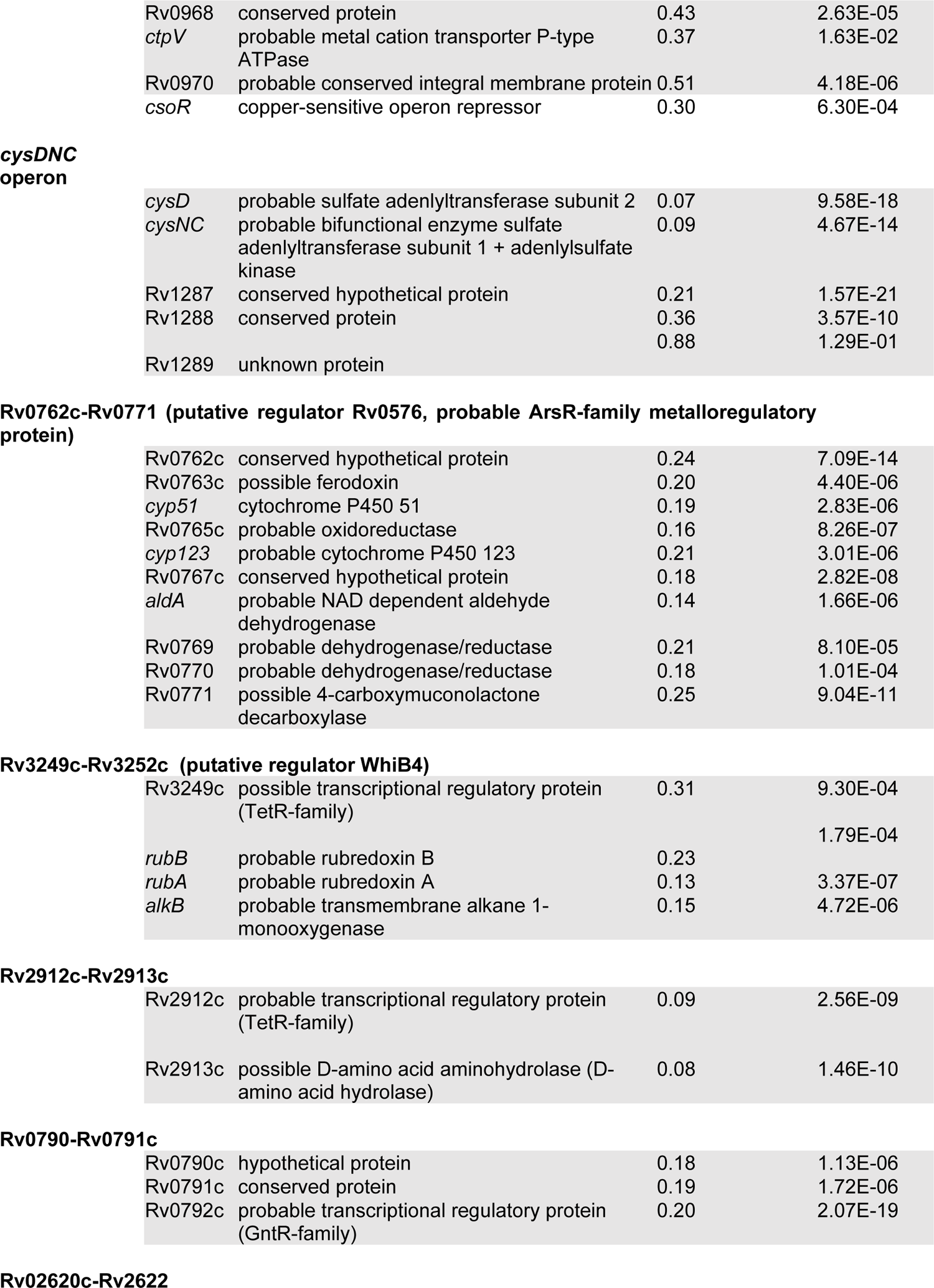
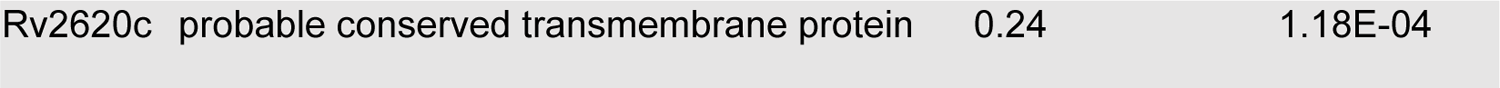
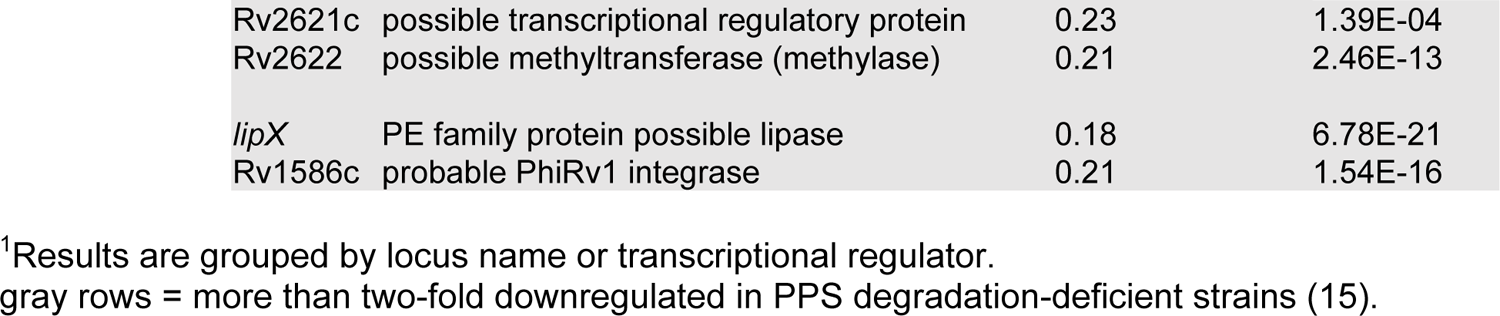
Downregulated operons and regulons in WT *M. tuberculosis* treated with pHBA relative to untreated bacteria.

**Table 3.**
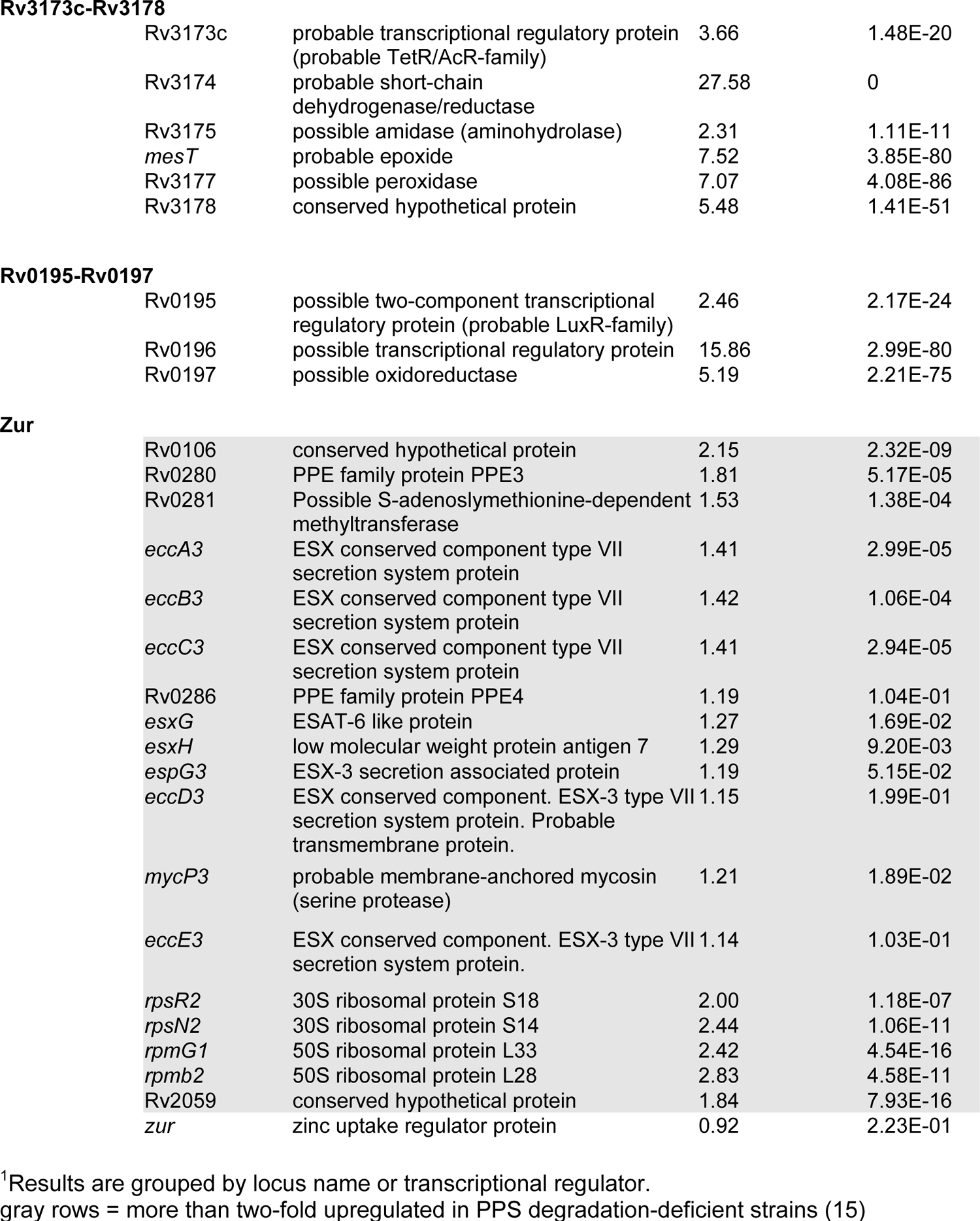
Upregulated operons and regulons in WT *M. tuberculosis* treated with *p*HBA relative to untreated.

**Rv3173c-Rv3178**
|  |  |  |  |
| --- | --- | --- | --- |
| Rv3173c | probable transcriptional regulatory protein (probable TetR/AcR-family) | 3.66 | 1.48E-20 |
| Rv3174 | probable short-chain dehydrogenase/reductase | 27.58 | 0 |
| Rv3175 | possible amidase (aminohydrolase) | 2.31 | 1.11E-11 |
| *mesT* | probable epoxide | 7.52 | 3.85E-80 |
| Rv3177 | possible peroxidase | 7.07 | 4.08E-86 |
| Rv3178 | conserved hypothetical protein | 5.48 | 1.41E-51 |
| **Rv0195-Rv0197** |  |  |  |
| Rv0195 | possible two-component transcriptional regulatory protein (probable LuxR-family) | 2.46 | 2.17E-24 |
| Rv0196 | possible transcriptional regulatory protein | 15.86 | 2.99E-80 |
| Rv0197 | possible oxidoreductase | 5.19 | 2.21E-75 |

|  |  |  |  |
| --- | --- | --- | --- |
| Rv0106 | conserved hypothetical protein | 2.15 | 2.32E-09 |
| Rv0280 | PPE family protein PPE3 | 1.81 | 5.17E-05 |
| Rv0281 | Possible S-adenosylmethionine-dependent methyltransferase | 1.53 | 1.38E-04 |
| *eccA3* | ESX conserved component type VII secretion system protein | 1.41 | 2.99E-05 |
| *eccB3* | ESX conserved component type VII secretion system protein | 1.42 | 1.06E-04 |
| *eccC3* | ESX conserved component type VII secretion system protein | 1.41 | 2.94E-05 |
| Rv0286 | PPE family protein PPE4 | 1.19 | 1.04E-01 |
| *esxG* | ESAT-6 like protein | 1.27 | 1.69E-02 |
| *esxH* | low molecular weight protein antigen 7 | 1.29 | 9.20E-03 |
| *espG3* | ESX-3 secretion associated protein | 1.19 | 5.15E-02 |
| *eccD3* | ESX conserved component. ESX-3 type VII secretion system protein. Probable transmembrane protein. | 1.15 | 1.99E-01 |
| *mycP3* | probable membrane-anchored mycosin (serine protease) | 1.21 | 1.89E-02 |
| *eccE3* | ESX conserved component. ESX-3 type VII secretion system protein. | 1.14 | 1.03E-01 |
| *rpsR2* | 30S ribosomal protein S18 | 2.00 | 1.18E-07 |
| *rpsN2* | 30S ribosomal protein S14 | 2.44 | 1.06E-11 |
| *rpmG1* | 50S ribosomal protein L33 | 2.42 | 4.54E-16 |
| *rpmb2* | 50S ribosomal protein L28 | 2.83 | 4.58E-11 |
| Rv2059 | conserved hypothetical protein | 1.84 | 7.93E-16 |
| *zur* | zinc uptake regulator protein | 0.92 | 2.23E-01 |

<sup>1</sup>Results are grouped by locus name or transcriptional regulator.
gray rows = more than two-fold upregulated in PPS degradation-deficient strains (15)

The Zur (<u>z</u>inc <u>u</u>ptake <u>r</u>egulator) regulon was upregulated in *p*HBA-treated *M. tuberculosis* (**Table 3**) and in PPS mutants (15). Zur is a zinc-responsive regulator that, unlike the Cu regulators, binds to the promoters of its regulon in the presence of excess zinc (21). Thus, Zur regulon genes are induced in low zinc and are therefore implicated in zinc uptake (21). The ESAT-6 cluster 3 (ESX-3) genes in the Zur regulon are also regulated by IdeR (iron <u>d</u>ependent <u>r</u>epressor) (22). However, no other IdeR-dependent genes were differentially expressed between untreated and *p*HBA-treated *M. tuberculosis*, similar to what we observed in PPS mutant *M. tuberculosis* strains (15).

In addition to the Zur regulon, several genes were induced in *p*HBA-treated *M. tuberculosis* that are not induced in PPS mutants (**Table 3**). Some of these genes are upregulated in hypoxia or after damage to the mycomembrane (23, 24). It is possible these genes are specifically induced in response to extracellular aldehyde exposure rather than endogenously-produced aldehyde as is found in PPS mutants. Alternatively, the amount of *p*HBA we used, which is above physiologic levels, induced a transcriptional response that would not be achieved by what are likely much lower concentrations of *p*HBA found in PPS mutants.

Given that *p*HBA-treated *M. tuberculosis* and PPS mutant strains are hypersensitive to Cu, and that the RicR regulon is responsible for conferring robust *in vitro* Cu resistance and virulence in mice (12), we tested if the observed downregulation of the RicR regulon resulted in lower protein levels of a gene product required for robust Cu resistance (12, 16). We compared levels of MmcO in *M. tuberculosis* strains lacking *mpa* and in WT *M. tuberculosis* treated with *p*HBA by immunoblotting. In the absence added Cu, *p*HBA treatment alone resulted in low MmcO levels in all strains (**Fig. 2**, lanes 1, 4 and 7). Cu treatment of the parental WT strain increased MmcO levels as previously reported (12) (**Fig. 2**, lane 2), whereas the addition of *p*HBA reduced levels of MmcO (**Fig. 2**, lane 3). The *mpa* strain showed low levels of MmcO, regardless of the addition of Cu (**Fig. 2**, lane 4-6) while the *mpa log* double mutant strain showed a similar pattern of MmcO levels as the WT strain (**Fig. 2**, lanes 2 v. 8, and lanes 3 v. 9). These results corroborate that the observed transcriptional changes in *p*HBA-treated WT and PPS mutant bacteria are due to the presence of one or more aldehydes.

**Fig. 2.**
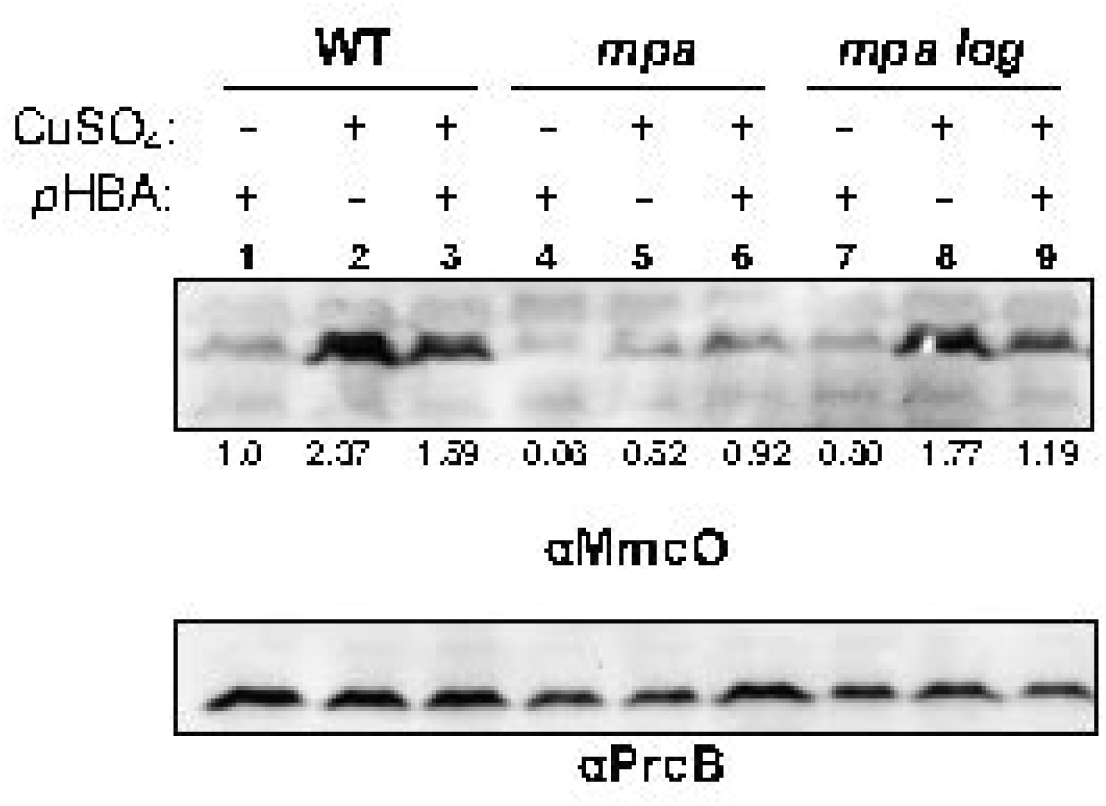
MmcO levels are reduced in a mutant defective for proteasomal degradation and in *p*HBA-treated *M. tuberculosis*. *M. tuberculosis* cultures grown in minimal Sauton media were treated with or without 1.2 mM *p*HBA 4 hours prior to treatment with 50 µM CuSO_4_. 24 hours after CuSO_4_ treatment, whole cell lysates were collected and prepared for immunoblotting using polyclonal rabbit antibodies raised against MmcO. Normalized MmcO levels (to PrcB levels) relative to MmcO level in WT treated with *p*HBA only (lane 1) are indicated under anti-MmcO blot. Antibodies against PrcB were used as a loading control. Data are representative of three independent experiments.

### *p*HBA alters the function of the Cu metallothionein MymT

*M. tuberculosis* encodes a single metallothionein, MymT, which is a low molecular weight, cysteine-rich protein that binds Cu (18). By binding specifically to reduced Cu [Cu(I)], metallothioneins protect cells from the potentially harmful effects of free Cu (25). Given that aldehydes can form adducts with cysteines in proteins and disable their function (26), we hypothesized *p*HBA could directly disrupt cysteines in MymT, preventing its ability to bind Cu. Cu-metallothioneins like MymT uniquely form solvent-shielded Cu(I)-thiolate cores that luminesce when excited by ultra-violet (UV) light (18, 27). We first monitored the formation of MymT Cu-thiolate cores in WT *M. tuberculosis* by fractionating whole-cell lysates of bacteria incubated with Cu and measuring luminescence emitted from each fraction after excitation by UV light. A sharp peak of luminescence was observed in the fractionation profile of WT *M. tuberculosis* that could be attributed to MymT, given that this peak was absent from lysates of a Δ*mymT::hyg* (“*mymT “)* strain (**Fig. 3A**).

**Fig. 3.**
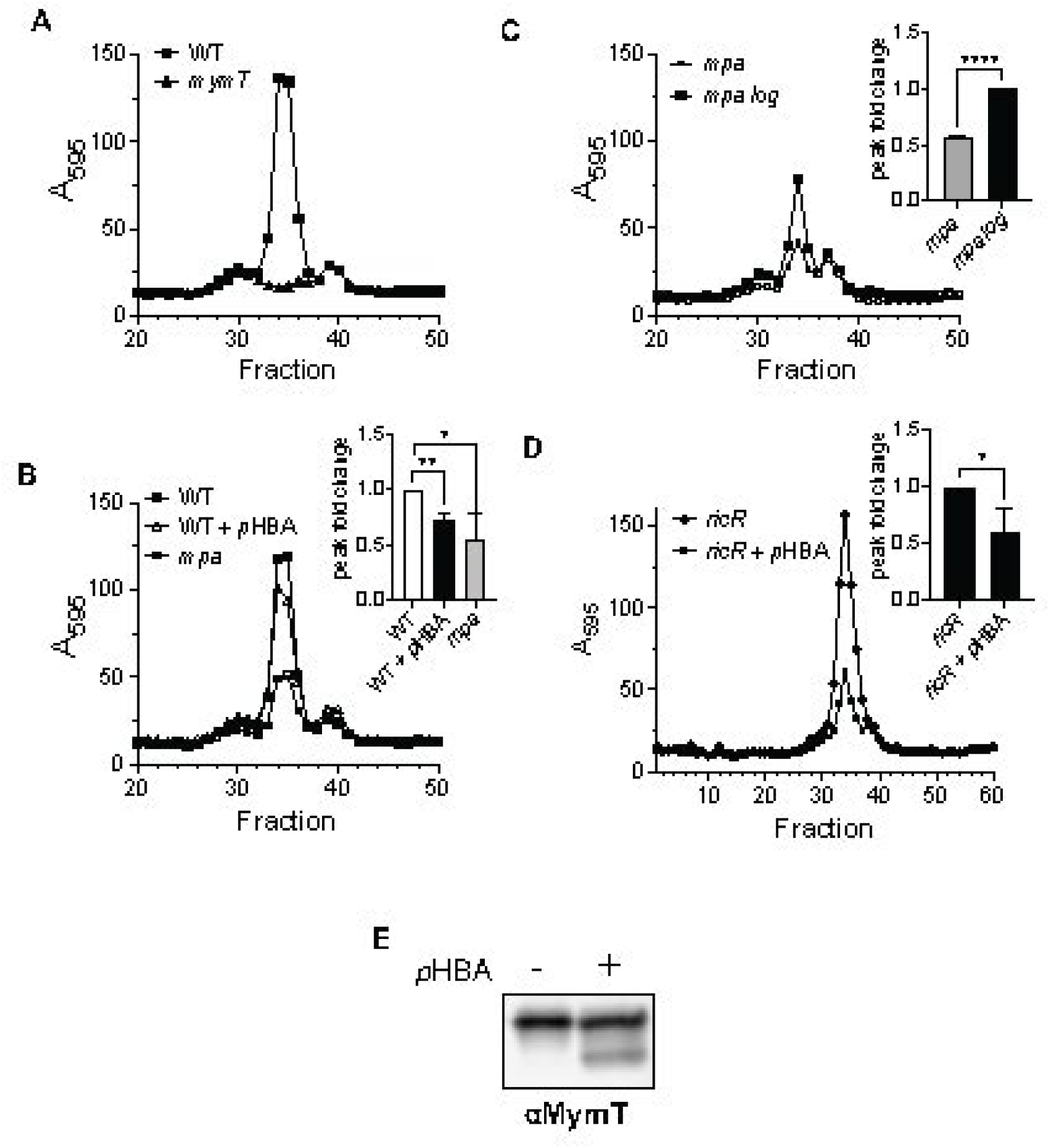
MymT binds less copper in the presence of aldehyde. MymT Cu(I)-thiolate luminescence in a PPS mutant and in *p*HBA-treated *M. tuberculosis* (excitation = 280 nm, emission = 595 nm, cutoff = 325 nm). **(A)** WT *M. tuberculosis* lysates (solid line, squares) exhibit a peak in luminescence in fraction 34, which is abolished in lysates of a *mymT* mutant (dashed line, triangles). **(B)** Fractionated lysates from WT, WT treated with *p*HBA, and *mpa* mutant *M. tuberculosis* strains. Inset is quantification of peak fold change for the three strains or conditions across three independent experiments. **(C)** Fractionated lysates from *mpa* and *mpa log* mutant strains. Inset: quantification of peak fold change between the two strains and two independent experiments. **(D)** Fractionated lysates from *a ricR* mutant treated with or without *p*HBA. Inset is quantification of peak fold change for the two conditions across three independent experiments. **(E)** Immunoblot for MymT fractions corresponding to the peak collected from **(D)**. All quantifications were analyzed for significance using an unpaired t-test with *P* < 0.05 (*) and *P* < 0.01 (**).

We next tested if MymT luminescence was reduced in an *mpa* mutant compared to in the parental strain. The MymT luminescence peak in the *mpa* strain was significantly lower (**Fig. 3B**). We also found that MymT luminescence could be reduced in WT *M. tuberculosis* in the presence of exogenously-added *p*HBA (**Fig. 3B**). Furthermore, the disruption of *log* in the *mpa* mutant restored the MymT luminescence peak (**Fig. 3C**).

A caveat to the experiments in Fig. 3B and C is that given *p*HBA-treated WT *M. tuberculosis* and untreated *mpa* strains have repressed RicR regulon expression, we could not conclude that the reduced luminescence of MymT was simply due to a reduction in its protein levels versus the disruption of its cysteines. To control for a difference in MymT levels, we repeated the experiments with a Δ*ricR::hyg* (“*ricR*”) strain that produces constitutively high levels of MymT (**Table 1**) (15). *p*HBA treatment of the *ricR* mutant resulted in a reduction of the MymT luminescence peak compared to the untreated culture (**Fig. 3D**). Importantly, we found MymT levels were equivalent between *p*HBA-treated and untreated cultures by immunoblotting for MymT in the fractions containing the luminescent peaks (**Fig. 3E**). Interestingly, we also consistently detected a smaller species of MymT in the *p*HBA-treated strain lysates. It is possible that this smaller species resulted from a covalent interaction of *p*HBA with MymT, for example, with the Cu-coordinating cysteines. As a result, it is possible the aldehyde-modified MymT not only could not bind Cu but also adopted a conformation that caused it to migrate through SDS-PAGE gels more quickly. Alternatively, modified MymT might adopt a conformation that exposes it to a peptidase resulting in the release of smaller peptides. Collectively, our data indicate that the reduction of MymT luminescence in a PPS degradation-deficient strain or after *p*HBA treatment was due to a combination of reduced MymT levels and disabled MymT Cu-binding. Furthermore, these results support a model whereby *p*HBA directly alters the ability of MymT to bind Cu.

### A metabolic aldehyde sensitizes *M. tuberculosis* to Cu

A recent hypothesis put forward by the Darwin and Stanley labs proposed host cell-derived aldehydes of metabolism contribute to bacterial control during infections (28). Macrophages undergo an increase in aerobic glycolysis, known as the Warburg Effect, following infection with *M. tuberculosis* and other pathogens (29-32). A by-product of aerobic glycolysis is MG, also known as pyruvaldehyde, which has been detected at millimolar concentrations in *M. tuberculosis*-infected mouse macrophages (33). Similar to *p*HBA, we found exogenously added MG could sensitize *M. tuberculosis* to Cu *in vitro* (**Fig. 4**). Thus, host-derived aldehydes have the potential to play a role in inactivating bacterial defenses against Cu toxicity during infections.

**Fig. 4.**
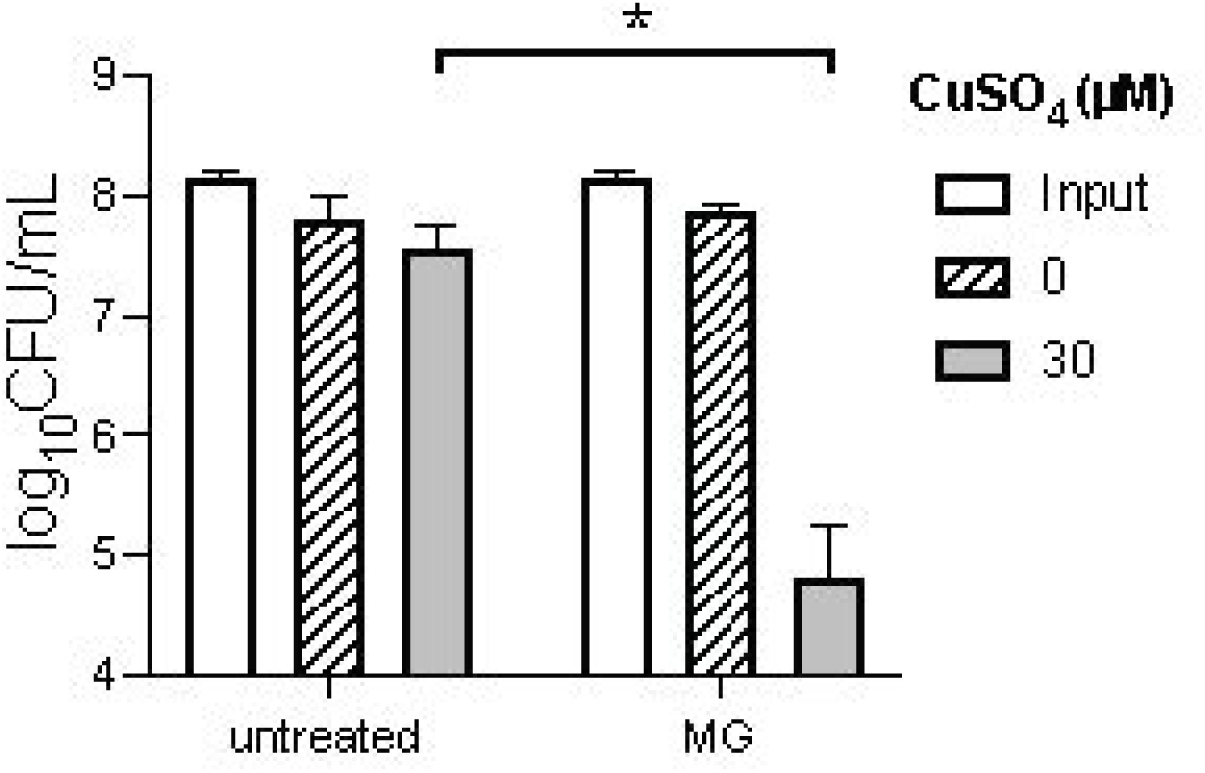
Methylglyoxal sensitizes *M. tuberculosis* to copper. Cu sensitivity assay was performed with WT *M. tuberculosis* treated with or without 360 µM MG and 30 µM of CuSO_4_. All quantifications were analyzed for significance using an unpaired t-test with *P* < 0.05 (*). Data are representative of two experiments, each done in technical triplicate.

## DISCUSSION

In this work, we sought to test if an aldehyde, *p*HBA, that accumulates in a PPS-deficient *M. tuberculosis* strain was responsible for its Cu-sensitive phenotype. Indeed, we found that disruption of *log*, which encodes a proteasome substrate that is the source accumulated *p*HBA, restored Cu resistance to a PPS mutant at levels comparable to the parental strain. We looked into other suppressors of NO sensitivity in PPS degradation-deficient strains and found that disruption of *glpK* also restored Cu resistance to WT levels. We found that the addition of *p*HBA to WT *M. tuberculosis* cultures was sufficient to sensitize bacteria to Cu and that this phenotype was likely due to the reduced the expression of genes needed for Cu resistance. We also showed that the metal-binding function of MymT was altered in the presence of *p*HBA. Thus, *p*HBA can affect both MymT levels and activity. Finally, we showed that an aldehyde produced by macrophages during infection, MG, could also sensitize *M. tuberculosis* to Cu *in vitro*. We therefore propose a model whereby aldehydes disable Cu-sensing by RicR, leading to the constitutive repression of Cu-resistance genes, and the disruption of Cu-binding by MymT; together, these effects result in Cu sensitivity (**Fig. 5**). In addition to the RicR regulon, aldehydes affected the expression of the CsoR operon, thus, it is also possible that interference with CsoR contributed to the increased Cu sensitivity of PPS mutants or aldehyde-treated WT *M. tuberculosis*.

**Fig. 5.**
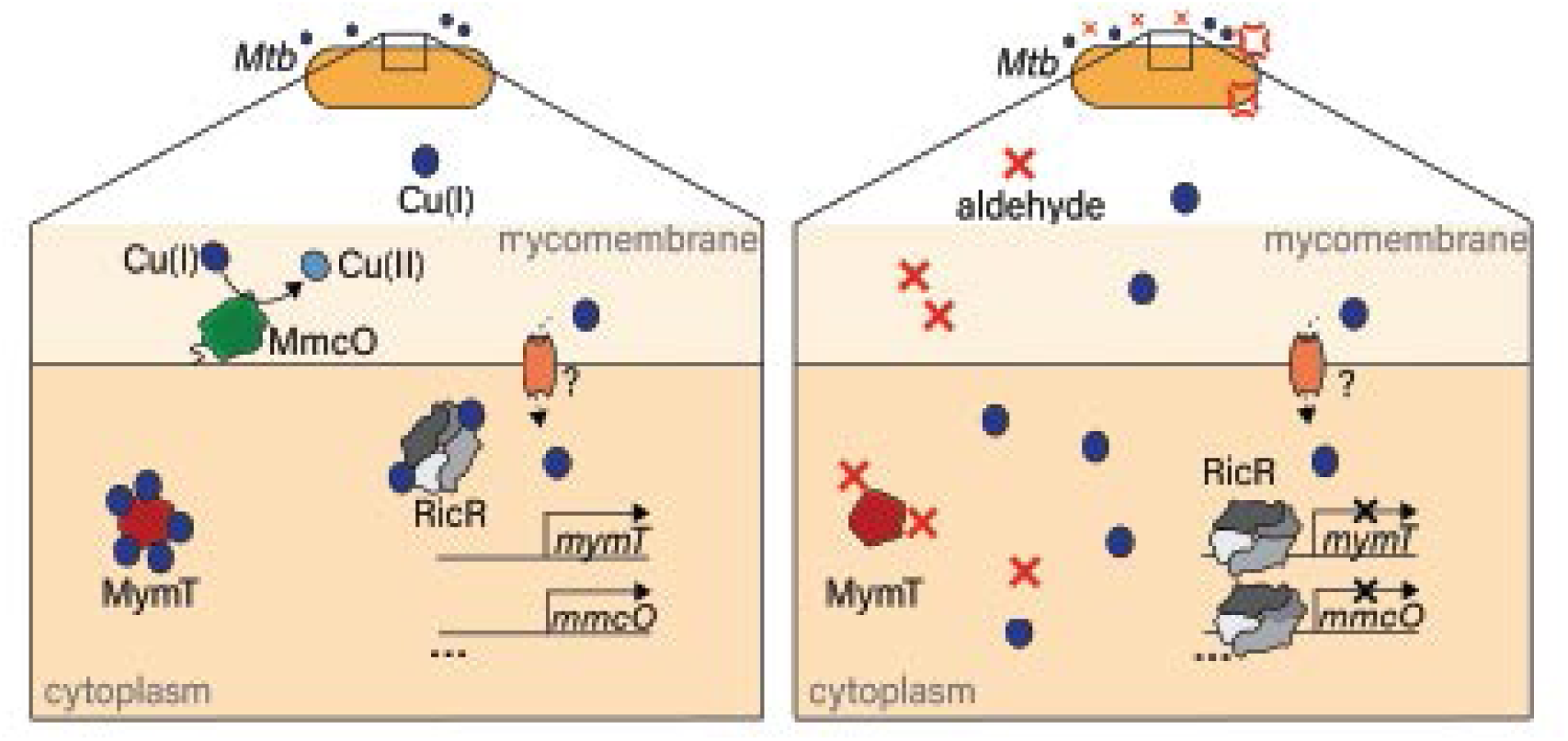
Model of aldehyde sensitization of *M. tuberculosis* to copper through disabling of the RicR regulon. Left panel: in the absence of aldehyde, Cu(I) enters the cytoplasmic space through unknown transporters. Cu prevents RicR from binding to DNA, allowing expression of the RicR regulon genes, some of which are implicated in countering Cu(I) toxicity in the bacteria. MymT sequesters Cu(I) to prevent it from damaging the cell, and MmcO is a periplasmic multicopper oxidase that oxidizes Cu(I) to its less toxic Cu(II). Right panel: in the presence of aldehyde, RicR cannot release from DNA even in the presence of Cu, leading to the constitutive repression of the RicR regulon and the observed Cu sensitivity of aldehyde-treated and PPS mutant *M. tuberculosis* strains. Additionally, MymT is unable to bind Cu in the presence of aldehyde.

While the focus of our study was on the aldehyde *p*HBA, the observation that disruption of *glpK* also suppresses both NO sensitivity (5) and Cu sensitivity (**Fig 1B**) could link the accumulation of additional aldehydes to these phenotypes. Recent studies by several groups found that *glpK*-deficient *M. tuberculosis* strains are prevalent in clinical isolates (34-36). To grow on glycerol, GlpK promotes the production of glyceraldehyde-3-phosphate, another aldehyde, as well as MG. Mutations in *glpK* are reversible and confer decreased susceptibility to antituberculosis drugs, suggesting phase variation of *glpK* occurs in response to antibiotic pressure on the bacteria. The Alland group proposed that the activation of a general stress response following a block in glycerol metabolism provides increased antibiotic tolerance (34). The Sassetti lab proposed that given that glycerol metabolism results in the production of MG, a block in this pathway could protect bacteria by lowering the aldehyde burden in the bacteria (35). The precise mechanism by which *glpK* mutations increase *M. tuberculosis* tolerance to drugs is unknown. Given the reactive nature of aldehydes and their ability to sensitize *M. tuberculosis* to molecules like NO and Cu, it is possible that reducing the production of endogenously-produced aldehydes can increase the intrinsic resistance of bacteria to antimicrobial drugs and other toxic molecules.

In addition to the Cu-responsive regulons, other downregulated genes were shared between *p*HBA-treated- and PPS mutant *M. tuberculosis* cultures. These included the *cysDNC operon*, which encodes a sulfate-activating enzyme complex that is implicated in virulence, oxidative stress, sulfate limitation, and sensing of exogenous cysteine (37); Rv0762c-Rv0771, which includes a gene for a putative aldehyde dehydrogenase (*aldA*) and is predicted to be regulated by Rv0576 (38), a probable ArsR-like metalloregulatory transcriptional repressor (39); and Rv3249c-Rv3252c that is predicted to be controlled by WhiB4, a redox-responsive, iron-sulfur cluster-containing transcriptional repressor (40). Rv3249c-Rv3252c encodes the rubredoxins RubA and RubB and a putative monooxygenase AlkB (**Table 2**, gray rows). Many of these genes are regulated by metal-sensing or metal-binding proteins including the ArsR-like repressors and WhiB4 that use cysteines to coordinate their respective metals (39, 40). Notably, the IdeR iron regulator, which does not use cysteine to coordinate its metal (22), was not dysregulated in *p*HBA-treated *M. tuberculosis* or in PPS mutants (15). Because aldehydes are reactive molecules that can form adducts with thiols in proteins, we propose that cysteine-dependent metal coordinating proteins are particularly sensitive to aldehyde exposure.

There have been other studies that have investigated the Cu response in *M. tuberculosis*. Most notably, a recent study by the Glickman lab identified an integrated system involving Rip1 protease and the PdtaS/R two-component system in *M. tuberculosis* that senses and mediates resistance to Cu and NO (41). Although a *rip1* mutant is highly sensitive to Cu, expression of the RicR and CsoR regulons appear to be unaffected. The NO sensitivity of a *rip1* mutant is attributed to a block in chalkophore biosynthesis. Chalkophores bind Cu with high affinity (41-43), but the restoration of expression of genes involved in their synthesis does not restore Cu resistance to a *rip1* mutant. While it remains to be determined how Cu and NO resistance is conferred by Rip1, it is unlikely that aldehydes are involved given that known Cu-resistance genes are not repressed in a *rip1* mutant. Importantly, these data suggest there are additional ways for *M. tuberculosis* to be sensitized to toxic molecules like NO and Cu.

While it is established Cu and NO are used by immune cells in defense against *M. tuberculosis*, an anti-microbial role for aldehydes *in vivo* has yet to be determined. It was recently proposed that aldehydes may be an underappreciated defense against invading pathogens (28). In macrophages, aldehydes are produced at low levels during cellular metabolism (44, 45), but a shift to aerobic glycolysis following infection with *M. tuberculosis* leads to an increase of aldehydes such as glyceraldehyde-3-phosphate and MG (29-31, 33, 46-48). Induction of aerobic glycolysis plays a role in infection control, given that the inhibition of this pathway in mouse macrophages leads to loss of some IFN-γ-dependent control of *M. tuberculosis* growth (29). Thus, aldehydes produced during this shift to glycolysis might contribute antibacterial activity for the host.

Our data represent the first evidence that aldehydes can sensitize *M. tuberculosis* to Cu. Importantly, despite the well-established evidence of their toxicity, the antibacterial mechanisms of aldehydes are relatively uncharacterized; thus, our study contributes new insight into how aldehydes can target and inactivate bacterial systems.

## MATERIALS AND METHODS

### Bacterial strains, growth conditions, plasmids, and primers

The bacterial strains, plasmids, and primers used in this work are listed in Table 1. *M. tuberculosis* strains were grown in 7H9 liquid broth (Difco) supplemented with 0.2% glycerol, 0.05% Tween 80, 0.5% bovine serum albumin (BSA), 0.2% dextrose, and 0.085% sodium chloride (ADN) (referred to as “7H9c” from here on) or Sauton minimal medium (3.7 mM potassium phosphate, monobasic; 2.4 mM magnesium sulfate; 30 mM L-asparagine; 3.5 mM zinc sulfate; 9.5 mM citric acid; 6.0% glycerol; 0.005% ferric ammonium citrate; 0.05% Tween-80). Cultures were grown at 37°C without agitation in vented flasks (Corning). For *M. tuberculosis* growth on solid medium, Middlebrook 7H11 agar (Difco and Remel) was supplemented with Middlebrook OADC (oleic acid, albumin, dextrose, and catalase; BBL). *M. tuberculosis* strains were grown in 50 µg/ml kanamycin, 50 µg/ml hygromycin when necessary.

For CuSO_4_ solutions: stock solutions were made by dissolving the appropriate amount of CuSO_4_ powder (Fisher Scientific) in water and filter-sterilized using a 0.45 µ filter. For aldehyde solutions: 50 mM *p*HBA was made with *p*HBA powder dissolved in water and filter-sterilized using a 0.45 µ filter. MG (5.5 M) was diluted in sterile water just before use. Both aldehydes were purchased from Sigma-Aldrich, Inc.

### RNA-Seq

*M. tuberculosis* cultures were grown in 7H9c media and treated with 1.2 mM *p*HBA at an optical density at 580 nm (OD_580_) of 1.0 (early stationary phase) or left untreated. 24 hours later, RNA was purified as previously described (15). RNA was isolated from three biological replicate cultures. Library preparation, and Illumina HiSeq Sequencing were performed by GENEWIZ, LLC. Sequence reads were mapped to the *M. tuberculosis* H37Rv genome sequence (RefSeq identifier GCF_000195955.2) using bwa v0.7.17 (49) and sorted using samtools v1.9 (50). An average of 97.8% of reads were mapped to the reference genome, indicating high quality of samples. Given a locus of interest, *socAB*, was not included in the assembly annotation from RefSeq, we added it to the annotation files at genomic coordinates NC_000962.3:1,933,937-1,934,497. Using the alignment files generated for each sample, the *featureCounts* command in Subread v2.0.1 (51) was used to count the reads mapping to each gene in the reference. Read counts per gene and sample (feature counts) were loaded into R v4.2.0 (52) for further analysis using the package DESeq2 v1.36.0 (53). DESeq2’s normalization function was used to normalize the counts to make expression levels more comparable between the different samples. To compare gene expression changes between *p*HBA-treated and untreated samples, the Wald test was used to generate *P*-values and log_2_-fold changes. Genes with an adjusted *P*-value of <0.05 and absolute log_2_-fold change of >2.0 when comparing *p*HBA-treated to untreated *M. tuberculosis* were considered differentially-expressed genes (**Table S1**). Raw sequencing data files are available in a PATRIC public workspace: (https://www.bv-brc.org/workspace/ginalimon@bvbrc/Limon_Darwin_RNA-Seq).

### Cu-sensitivity assay

Cu-sensitivity assays were performed as previously described (12, 15). Briefly, *M. tuberculosis* strains were grown in 7H9c to an OD_580_ = 0.5-1.0. Bacteria were washed once with Cu-free Sauton media and collected using a low-speed centrifugation (150 x g) to remove clumped cells. Supernatants containing mostly unclumped bacteria were diluted to OD_580_ = 0.08 in Sauton medium. 194 µl of this diluted culture were transferred to wells of a 96-well plate; 6 µl of the appropriate stock concentration of CuSO_4_ or aldehyde or both were added to the desired final concentration. Plates were incubated at 37ºC for 10 days, after which cultures were diluted and inoculated onto 7H11 Y-agar plates. Plates were incubated for 14-21 days before enumerating CFU. As previously reported (15), we used a range of CuSO_4_ concentrations due to variability of Cu-sensitivity between experiments. Each experiment was done at least twice, each with technical triplicates.

### *M. tuberculosis* lysate preparation for immunoblotting

For all blots, bacteria were grown in Sauton media with no added Cu. For MmcO blots, at OD_580_ = 0.5-1.0, cultures were treated with 1.2 mM *p*HBA for 4 hours before treatment with 50 µM CuSO_4_. Bacteria were harvested 24 hours after the addition of CuSO_4_. Bacterial densities were measured and equivalent cell numbers were collected, based on the OD_580_ of the cultures. For example, an “OD_580_ equivalent of 1” indicates the OD_580_ of a 1 ml culture is 1.0. For most assays, 5 OD_580_ units were collected and washed once with phosphate-buffered saline (DPBS, Corning, Inc with 0.05% Tween-80) to remove BSA in 7H9c media. Bacterial pellets were then resuspended in 300 µl TE buffer (100 mM Tris-Cl, 1 mM EDTA pH 8.0), and transferred to bead-beating tubes with 200 µl zirconia beads; tubes were beaten for 30 seconds three times, with icing for 30 seconds in between in a mini bead beater (all materials from Bio-Spec). 150 µl of lysate was transferred into new tubes with 50 µl 4× SDS sample buffer (250 mM Tris pH 6.8, 2% SDS, 20% β-mercaptoethanol or BME, 40% glycerol, 1% bromophenol blue) and boiled at 100ºC for 10 minutes. Proteins were separated by sodium dodecyl sulfate polyacrylamide gel electrophoresis (SDS-PAGE) and transferred onto nitrocellulose membranes. Membranes were blocked in 3% milk or BSA prior to incubation with polyclonal rabbit antibodies as indicated in the figure legends. For loading controls, the same membranes were stripped with 0.2 N NaOH as described elsewhere (54) and re-blocked before incubation with antibodies to *M. smegmatis* PrcB (55).

### Construction of a Δ*ricR*::*hyg* mutant

We made an *M. tuberculosis ricR* deletion mutant strain using a previously described method (15). Briefly, pYUB854 (56) was used to clone sequences encompassing ∼700 bp upstream (5’) and ∼700 downstream (3’) of the *ricR* gene. The 5’ and 3’ sequences, including the start and stop codons, respectively, were cloned to flank the hygromycin resistance cassette in pYUB854. The plasmid was digested with PacI, and approximately 1 µg of linearized, gel-purified DNA was used for electroporation into *M. tuberculosis. M. tuberculosis* strains were grown to an OD of ∼0.4 to 1, washed, and resuspended in 10% glycerol to make electrocompetent cells as described in detail in (57). Bacteria were inoculated onto 7H11 agar with 50 µg/ml hygromycin as needed; a no-DNA control electroporation was done to control for spontaneously antibiotic-resistant mutants. Two weeks after plating, colonies were picked and inoculated into 200 µl 7H9c with antibiotics, and then inoculated into 5 ml cultures for further analysis. Mutants were confirmed by PCR and sequence analysis.

### MymT luminescence from *M. tuberculosis* lysates

We adapted a previously reported protocol for measuring Cu(I)-thiolate core luminescence for use on filtered *M. tuberculosis* lysates (18). *M. tuberculosis* cultures were grown in Sauton media to an OD_580_ = 0.3-0.5 and treated with *p*HBA to a final concentration of 1.2 mM as needed, and 50 µM CuSO_4_ four hours later. 24 hours after Cu addition, 12 OD_580_ equivalent cell numbers were harvested by centrifugation and washed twice with buffer (10 mM HEPES, 150 mM NaCl pH 7.4). Bacteria were resuspended in 700 µl of the same buffer and lysed by bead beating as described for preparing lysates for immunoblotting. Lysates were then centrifuged for 7.5 min at 20,000 *g* and the supernatants were passed through a 0.2 µ spin filter twice before application onto a Superose-6 10/300 GL column (Cytiva). Fractions were transferred to a UV grade 96-well plate (Corning) and luminescence was measured with excitation at 280 nm, emission 595 nm, and a cutoff of 325 nm.

For immunoblotting proteins in fractionated lysates: 200 µl of fractions corresponding to the three at the peak fluorescence collected using method above were frozen at −20ºC before being concentrated in a 0.5 ml centrifugal filter (Amicon). Samples were boiled in SDS sample buffer for 10 minutes before separation on 15% SDS-PAGE gels and transferred onto nitrocellulose membranes. Polyclonal MymT antibodies used for immunoblotting were a kind gift from Ben Gold and Carl Nathan (18).

## ACKNOWLEDGEMENTS

We thank S. Becker, S.A. Stanley and P. Tran for reviewing a draft version of this manuscript. We thank X. Shi for making MHD799. This work would not have been possible without preliminary experiments by M. Samanovic-Golden, the technical expertise of J. Ilmain (V. Torres Lab), or advice from S. Kahne. We thank J. Belasco, K. Cadwell, and M. Pacold for helpful suggestions. This work was supported by NIH grant AI153197 awarded to K.H.D and S.A.S. We thank the Office of Science and Research High-Containment Laboratories at NYU Grossman School of Medicine for their support in the completion of this research.

